# Extracellular vesicles alter trophoblast function in pregnancies complicated by COVID-19

**DOI:** 10.1101/2024.02.17.580824

**Authors:** Thea N. Golden, Sneha Mani, Rebecca L. Linn, Rita Leite, Natalie A. Trigg, Annette Wilson, Lauren Anton, Monica Mainigi, Colin C. Conine, Brett A. Kaufman, Jerome F. Strauss, Samuel Parry, Rebecca A. Simmons

## Abstract

Severe acute respiratory syndrome coronavirus 2 (SARS-CoV-2) infection and resulting coronavirus disease (COVID-19) causes placental dysfunction, which increases the risk of adverse pregnancy outcomes. While abnormal placental pathology resulting from COVID-19 is common, direct infection of the placenta is rare. This suggests that pathophysiology associated with maternal COVID-19, rather than direct placental infection, is responsible for placental dysfunction and alteration of the placental transcriptome. We hypothesized that maternal circulating extracellular vesicles (EVs), altered by COVID-19 during pregnancy, contribute to placental dysfunction. To examine this hypothesis, we characterized maternal circulating EVs from pregnancies complicated by COVID-19 and tested their effects on trophoblast cell physiology *in vitro*. We found that the gestational timing of COVID-19 is a major determinant of circulating EV function and cargo. *In vitro* trophoblast exposure to EVs isolated from patients with an active infection at the time of delivery, but not EVs isolated from Controls, altered key trophoblast functions including hormone production and invasion. Thus, circulating EVs from participants with an active infection, both symptomatic and asymptomatic cases, can disrupt vital trophoblast functions. EV cargo differed between participants with COVID-19 and Controls, which may contribute to the disruption of the placental transcriptome and morphology. Our findings show that COVID-19 can have effects throughout pregnancy on circulating EVs and circulating EVs are likely to participate in placental dysfunction induced by COVID-19.

## INTRODUCTION

Maternal SARS-CoV-2 infection and resulting coronavirus disease (COVID-19) is associated with an increased risk of pregnancy complications including preterm birth, hypertensive disorders of pregnancy, fetal growth restriction, and pregnancy loss (*1, 2*). Placental dysfunction is known to contribute to these complications, and placental pathology, including vasculopathies and inflammation, is frequently reported following an acute or even resolved infection during pregnancy (*3–6*). This suggests COVID-19 has a long-lasting effect on pregnancy by altering placenta function. Despite extensive reports of placental abnormalities following COVID-19, little is known about the underlying mechanisms contributing to placental dysfunction and the related subsequent pregnancy complications. Direct infection of the placenta is rare, which suggests that placental dysfunction is caused by the maternal response to SARS-CoV-2 infection (*6–8*).

Circulating extracellular vesicles (EVs) are altered by SARS-CoV-2 infection and contribute to COVID-19-induced organ damage (*9–13*). EVs are a means of cell-to-cell communication resulting from their ability to carry and transfer bioactive cargo that elicits signaling events in recipient cells. Compared to uninfected individuals, EV cargo composition in patients with COVID-19 is significantly different, eliciting downstream systemic effects such as coagulopathy (*14–17*) and inflammation (*10, 11, 14, 18*). For example, tissue factor protein abundance in EVs is increased in COVID-19 and correlates with inflammation and disease severity demonstrating the influence that circulating EVs have on the systemic response to COVID-19, thereby leading to organ dysfunction (*9, 12, 13*).

During normal pregnancy, the placenta releases EVs into the maternal circulation (*19, 20*). Placental-derived EVs promote maternal adaptation to support a healthy pregnancy including a shift in the maternal immune system to a tolerant state and the promotion of angiogenesis (*21, 22*). Placental-derived EVs also affect trophoblast function through autocrine and paracrine signaling. Trophoblasts are a specialized cell type of the placenta that are responsible for invasion into maternal tissue to anchor the placenta, vascular remodeling for adequate placental blood flow, nutrient transport, and hormone production for maternal and fetal signaling. EV signaling impairs normal trophoblast function which is thought to contribute to the placental dysfunction underlying pregnancy complications (*21, 23–26*).

Therefore, we hypothesized that COVID-19 alters circulating EV cargo which has a functional consequence in the placenta. Similar to previous studies, we found that COVID-19 during pregnancy induced marked placental histologic and transcriptomic changes that were dependent on the gestational timing of maternal SARS-CoV-2 infection. Importantly, we found that COVID-19 altered EV cargo, and trophoblast exposure to these EVs resulted in reduced trophoblast invasion and hormone production.

## RESULTS

### Pregnancy outcomes following active and resolved SARS-CoV-2 infection

Participants were enrolled at the time of delivery between July 2020-August 2022. Controls had no known SARS-CoV-2 infection during pregnancy. COVID-19 cases were divided based on the gestational timing of infection and either had a resolved infection that occurred in the 1^st^, 2^nd^, or 3^rd^ trimester (R1, R2, R3), or had an active infection at the time of delivery (AI) (Table 1). All patients admitted to the labor and delivery unit underwent PCR testing for SARS-CoV-2 at admission. There were no significant differences in maternal age between groups, but there were significantly more Black participants with an infection in the third trimester (resolved or active) compared to non-infected individuals (Controls) (R3 p=0.046, AI p=0.011). Participants with an infection in the third trimester, resolved or active, also had an earlier gestational age at delivery than Controls (R3 p<0.001, AI p=0.034) (Table 1). Not surprisingly, the AI group, which underwent universal screening for SARS-CoV-2 at admission for delivery, had a higher incidence of asymptomatic infection than those with resolved infection (p<0.0001).

**Table 1.** Subject demographics, pregnancy outcomes, and placenta pathology Participants enrolled in the COMET study formed five groups (controls, resolved infection in the 1^st^ trimester (R1), 2^nd^ trimester (R2), and 3^rd^ trimester (R3), and active infection (AI). Maternal demographics including maternal age, race, and gestational age (GA) at birth are reported. The severity of COVID-19 during their pregnancy, incidence of pregnancy complications (gestational hypertension (gHTN), chronic hypertension (cHTN), preeclampsia (PE), spontaneous and medically indicated preterm birth (PTB), intrauterine growth restriction (IUGR), and intrauterine demise (IUFD)) and placental pathology (maternal vascular malperfusion (MVM), fetal vascular malperfusion (FVM) and perivillous fibrin deposition) are reported * p<0.05 chi-squared test compared to Controls. ^+^ p<0.05 compared to other COVID-19 groups

| Demographics | Controls<br>(n=32) | R1<br>(n=15) | R2<br>(n=20) | R3<br>(n=22) | AI<br>(n=21) |
| --- | --- | --- | --- | --- | --- |
| Maternal age (years) | 21 – 42<br>(Mean: 31.8) | 25 - 43<br>(Mean: 31.7) | 18 – 39<br>(Mean: 29.4) | 21 – 42<br>(Mean: 30.5) | 20 – 40<br>(Mean: 29.4) |
| Race: White | 50% | 46.7% | 34% | 27.3% | 9.5%* |
| Race: Black | 41% | 46.7% | 55% | 68.2%* | 76.2%* |
| Race: Asian | 9.4% | 0% | 10% | 0% | 9.5% |
| Race: Other/Unknown | 0% | 6.7% | 0% | 4.5% | 4.8% |
| GA at delivery (weeks) | 37.3 – 41.1<br>(Mean: 39.3) | 35.1 – 40.3<br>(Mean: 39.1) | 33.7 – 41.3<br>(Mean: 38.4) | 32.3 – 39.7<br>(Mean: 37.5) * | 31 – 41<br>(Mean: 38.1) * |
| COVID severity | Controls<br>(n=32) | R1<br>(n=15) | R2<br>(n=20) | R3<br>(n=22) | AI<br>(n=21) |
| Symptomatic | N/A | 73.3% | 85% | 81.8% | 14.3% |
| Asymptomatic | N/A | 26.7% | 15% | 18.2% | 85.7%<sup>+</sup> |
| Hospitalized | N/A | 13.3% | 0% | 31.8% | 4.8% |
| Pregnancy outcomes | Controls<br>(n=32) | R1<br>(n=15) | R2<br>(n=20) | R3<br>(n=22) | AI<br>(n=21) |
| GHTN | 18.8% | 53.3%*<sup>+</sup> | 20% | 13.6% | 9.5% |
| CHTN | 3.1% | 0% | 5% | 13.6% | 4.8% |
| Preeclampsia | 0% | 0% | 5% | 18.2%* | 19%* |
| Spontaneous PTB | 0% | 0% | 20%* | 4.5% | 4.8% |
| Medically indicated PTB | 0% | 6.7% | 10% | 18.2%* | 19%* |
| IUGR | 6.3% | 13.3% | 0% | 9.1% | 4.8% |
| IUFD | 0% | 0% | 0% | 4.5% | 0% |
| Placenta Pathology | Control<br>(n=26) | R1<br>(n=9) | R2<br>(n=17) | R3<br>(n=19) | AI<br>(n=19) |
| MVM or FVM | 26.9% | 100%* | 71%* | 63%* | 82.6%* |
| MVM | 15% | 44%* | 47%* | 37% | 53%* |
| High grade MVM | 0% | 11% | 18%* | 16%* | 11% |
| FVM | 15% | 78%* | 59%* | 42%* | 47%* |
| >10% perivillous fibrin deposition | 0% | 11% | 18%* | 21%* | 5% |

Multiple studies have shown that SARS-CoV-2 infection during pregnancy is associated with adverse pregnancy outcomes. However, to date, no study has examined the relationship between the timing of infection and pregnancy complications in a single cohort. Our study is limited by relatively small numbers (n=15-32) in each participant group, but we found that the type of adverse outcome differed depending on the gestational timing of infection. The incidence of gestational hypertension (gHTN) was increased in R1 (p=0.016), whereas spontaneous preterm birth (SPTB) was increased in R2 (p=0.008) compared to Controls. Preeclampsia (PE) and medically indicated preterm birth (MPTB) were increased in R3 and AI compared to Controls (PE: R3 p=0.032, AI p= 0.010, MPTB: R3 p=0.005, AI p=0.010) (Table 1). In our cohort, there was no increase in intrauterine growth restriction (IUGR) or intrauterine fetal demise (IUFD) which have been reported in pregnancies complicated by COVID-19 (*27–29*).

### Placental pathology in COVID-19

Abnormal placental pathology is commonly reported in patients with active and resolved SARS-CoV-2 infections (*3*). However, there has yet to be a comprehensive assessment of placenta morphology following maternal infection at various gestational ages. Similar to previous reports (*3–6, 30, 31*), we found maternal and fetal vascular malperfusion (MVM & FVM) lesions were increased among patients with pregnancies complicated by COVID-19 (Table 1). Interestingly, high-grade MVM and perivillous fibrin deposition were increased in participants with a resolved infection that occurred in the second or third trimester, but not the first trimester or with an active infection. This suggests that it takes a substantial amount of time for the placenta to recover from the impact of COVID-19. Alternatively, it is possible that the first trimester placenta was more resilient. The lack of high-grade MVM and perivillous fibrin deposition in placenta collected from participants with an active infection suggests these lesions may take weeks to manifest.

### Timing of COVID-19 impacts the placental transcriptome

To gain insight into potential novel pathways that might be affected by COVID-19 during pregnancy, we performed RNA-seq on the placental transcriptome using biopsies from placenta collected at delivery in COVID-19 cases and controls. Not surprisingly, active infection was associated with significant differences in gene expression in the placenta. There were 72 upregulated genes and 384 downregulated genes in AI compared to Control placentas. (Figure 1, Supplemental Table 1). Gene expression was also altered in recovered infections, but the magnitude of change was smaller; 28, 2, and 56 differentially expressed genes (DEGs) comparing R1, R2, and R3 to Controls, respectively. It’s worth noting that in each of the 4 groups, there was differential expression of genes that regulate mitochondria activity, which implies a common dysfunctional pathway resulting from COVID-19.

**Figure 1.**
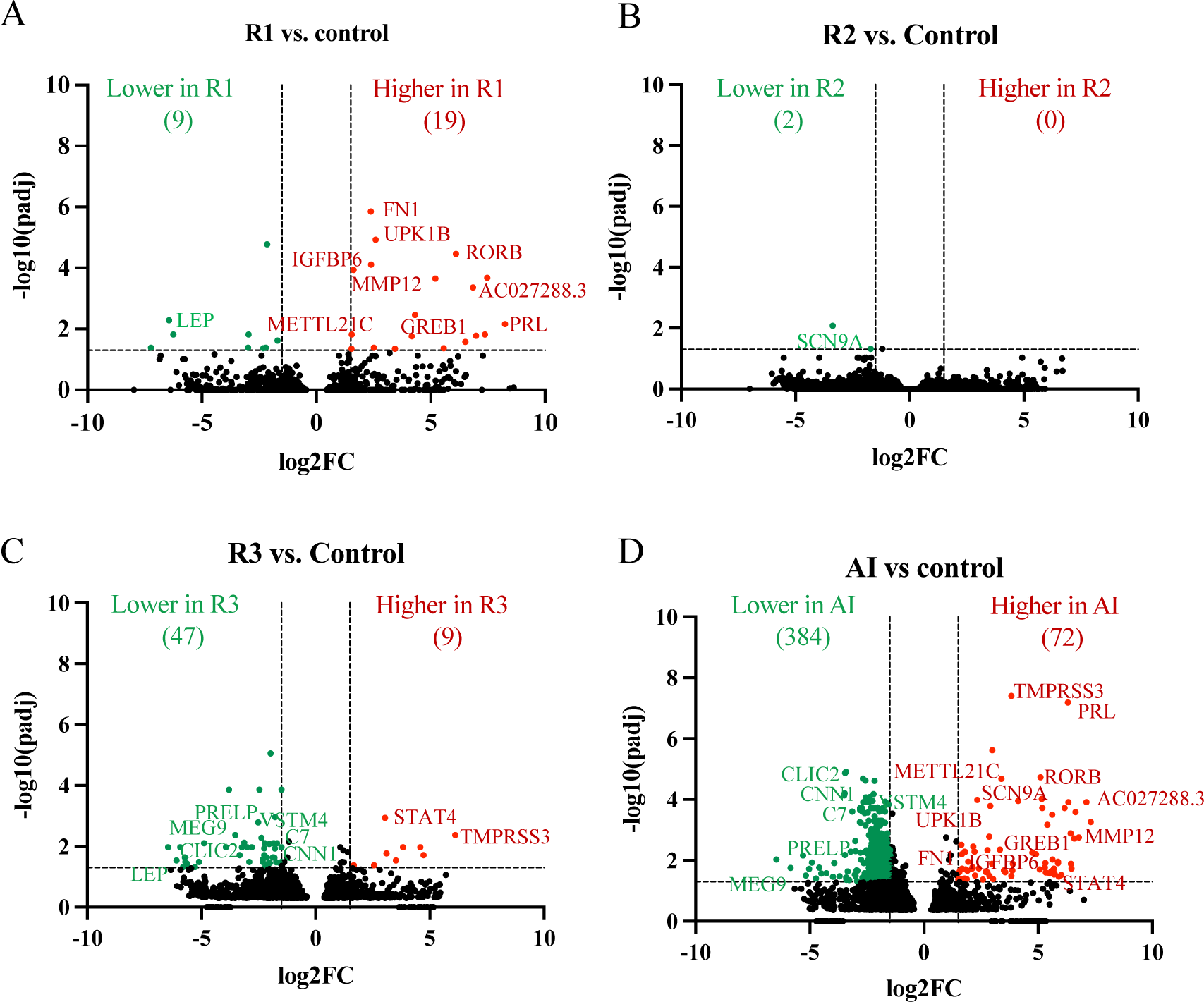
Differential gene expression in the placentas from patients with COVID-19 during pregnancy are represented by volcano plots. **(A-D)** The number and direction of differentially expressed genes between Controls (n=5) and resolved infections in the first trimester (R1) (A), second trimester (R2) (B), and third trimester (R3) (C), and active infection (AI) (D) is listed at the top of each graph. The gene name for those transcripts with differential expression in more than one COVID-19 group compared to Control is listed. (n=3-5/group)

In addition, several other genes were differentially expressed in one or more COVID-19 groups. There were several transcripts that were increased in placenta from R1 and AI groups compared to Controls. Several of these genes regulate fibrosis (*RORB*, *FN1*, *IGFB6*, *MMP12*, and *AC027288.3*) (*32–36*) suggesting that this pathway plays an important role in placenta pathology in COVID-19. Moreover, MMP12 regulates spiral artery remodeling and reduced MMP12 activity contributes to the development of preeclampsia (*37*). Similarly, increased expression of *FN1* slice variants containing Extra Domain A promotes inflammation via Toll-like receptor 4 (TLR4) activation (*38, 39*) and is associated with an increased risk of preeclampsia (*32, 38, 39*). Dysregulation of these pathways may contribute to the pathogenesis of preeclampsia in pregnant individuals with COVID-19. Finally, *GREB1* was upregulated in R1 and AI placenta compared to Controls. GREB1 interacts with the progesterone (P4) receptor to regulate P4 responsive genes (*40, 41*) and GREB1 promotes maternal tissue remodeling (*40*).

Several of the downregulated genes in AI compared to Controls were also downregulated in R3 placenta compared to Controls (*PRELP*, *MEG9*, *VSTM4*, *CLIC2*, *C7*, and *CNN1*). Interestingly, *CNN1*, which encodes calponin, is expressed by smooth muscle cells and expression changes are associated with spiral artery remodeling (*42, 43*). Further, SARS-CoV-2 infection is known to disrupt complement pathways (*44*) and the gene encoding complement C7 was downregulated in R3 and AI placenta compared to Controls. The complement system plays a dual role in pregnancy in that it protects the placenta from pathogen infection and participates in spiral artery remodeling (*45*). While the effect of decreased *C7* in the placenta is unknown, it may indicate that an imbalance in the complement system contributes to placental pathology observed in R3 and AI placentas. *STAT4*, which encodes the signal transducer and activator of transcription 4, is a key activator of immune regulating genes and was upregulated in R3 and AI placentas compared to Controls. STAT4 mediated pathways are disrupted in preeclampsia and circulating levels are elevated in patients with preeclampsia (*46, 47*). This differential gene expression was only seen in the placenta if the maternal infection was active or recently resolved, suggesting a long period of time is necessary to attenuate these pathways.

Statistically significant disruptions in canonical pathways, as determined by Ingenuity Pathway Analysis (IPA), include inflammation and fibrosis in the placentas from R1, R3, and AI compared to Controls (Supplemental Table 2). This unbiased approach based on differential gene expression supported our placental histopathology findings of inflammation and fibrosis in these placentas. Thus, despite the long period of time following the resolution of maternal SARS-CoV-2 infection, genes that regulate fibrosis and inflammation were altered, and pathological evidence of fibrosis and inflammation were apparent in the placenta regardless of the timing of infection.

We also used IPA to predict transcriptional regulators of genes differentially expressed in the placenta from pregnancies complicated by COVID-19. Interestingly, genes that encode for growth factors (IGF1, *IGF2*, *FGF3, FGF19, TGFB1*), immune-regulating proteins (*JUN, TNF, IL1B, IL13, IL6, IL10, IL4, IFNG*) and hormone-regulating proteins (*PRLH, LEPR, ESR1, and PGR*) were the top predicted transcriptional regulators (Supplemental Table 3).

### Sustained effects on circulating EVs following COVID-19 in early pregnancy

As discussed above, SARS-CoV-2 rarely infects the placenta, implying a distal signal. We hypothesized that maternal circulating EVs play a role in mediating placental dysfunction associated with COVID-19. Therefore, we characterized EVs isolated from maternal plasma collected at delivery to determine if COVID-19 altered the EV profile. We isolated large (LEV) and small (SEV) EVs as they carry distinct cargo with distinct functional effects. We confirmed the presence of large and small EVs in isolated particles by electron microscopy (Supplemental Figure 1A). Large and small EVs had the expected size distribution of EVs (Supplemental Figure 1B) and abundant expression of the EV-related tetraspanin CD9 (Supplemental Figure 1C).

EV characteristics, including concentration and size distribution, revealed long-lasting alterations in patients with a resolved infection. The diameter of small EVs was significantly increased in resolved infections compared to Controls (108.7nm vs. 117.2nm, p=0.023). However, the difference in concentration was not significant (2.15×10^8^ vs. 1.56 ×10^8^ EVs/uL plasma, p=0.074). When resolved infections were categorized by timing during gestation, we found that small EVs isolated from R2 participants had an increased diameter and reduced concentration, but small EVs isolated from R1 and R3 participants were not different from Controls (Figure 2 D&E). The diameter of large EVs isolated from R2 patients was decreased but there was no change in their number (Figure 2 A&B). COVID-19 in the second trimester was uniquely associated with alterations in circulating EV concentration and size at the time of delivery.

**Figure 2.**
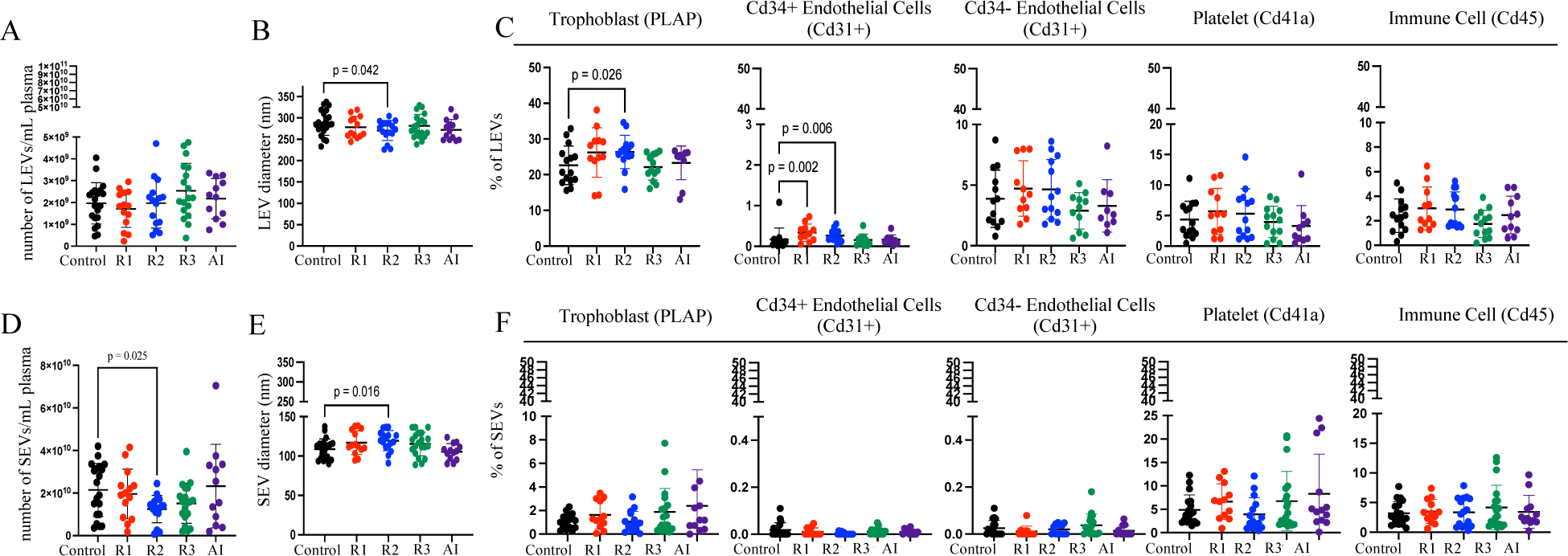
Circulating EVs were persistently altered in participants who experienced COVID-19 in the second trimester. **(A)** The number of LEVs in the circulation at the time of delivery (n=14-22/group). **(B)** The diameter of LEVs in the circulation (n=14-22/group). **(C)** Relative frequency of LEVs derived from trophoblasts, endothelial cells, platelets, and immune cells (n=11-16). **(D)** The number of small EVs in the circulation at the time of delivery (n=14-22/group). **(E)** The diameter of small EVs in the circulation (n=14-22/group). **(F)** The relative frequency of small EVs derived from endothelial cells, platelets, immune cells, and trophoblasts (n=13-20/group). All data are presented as mean ± SD. All analyses were performed by one-way ANOVA or the Kruskal-Wallis test, followed by post-hoc tests. Comparisons were made between Controls and resolved infection in the first trimester (R1), second trimester (R2), third trimester (R3), and active infection (AI).

### Altered cell of origin of EVs in patients with COVID-19 in early pregnancy

Characterizing the source of circulating EVs provides biological information about the tissue and cell-type of origin and its functional state. We used flow cytometry to detect cell-specific vesicle membrane protein expression and identified the relative contribution of each EV tissue-cellular source. We identified EVs that originated from maternal endothelial cells (CD31+ CD34-), fetal endothelial cells (CD31+ CD34+), platelets (CD41a+), immune cells (CD45+), and trophoblasts (PLAP+) (Figure 2 C&F).

Trophoblast-derived PLAP+ EVs comprised the largest proportion of circulating large EVs (Figure 2C). The percentage of PLAP+ EVs was increased in the circulation of R2 compared to Controls suggesting the placenta secreted more large EVs into circulation. Interestingly, we found a subset of endothelial-derived EVs that also express CD34, suggesting that these EVs originated from fetal endothelial cells (*48*). Fetal endothelial cell-derived large EVs (CD34+ CD31+) were also elevated in R1 and R2 compared to Controls. The percentage of small EVs from the placenta was not altered by COVID-19 (Figure 2F). In fact, there was no difference in the percentage of small EVs from any cell type measured. This suggests that placenta-derived large, but not small EVs, were altered by COVID-19 in early pregnancy.

### Circulating EVs from COVID-19 pregnant patients alter trophoblast function *in vitro*

The placenta is made up of three main functional cell types; 1) syncytiotrophoblast cells, which are responsible for nutrient transport and hormone production; 2) cytotrophoblast cells, the replicating precursors of the syncytiotrophoblast; and 3) extravillous trophoblasts (EVT), which invade deep into the uterus to anchor the placenta and enable blood and nutrient flow to the fetus. EV signaling is known to influence trophoblast function (*49*). Therefore, we tested the capacity of circulating EVs isolated from participants with an active infection or Controls to alter the function of trophoblast cell types. We focused our *in vitro* experiments on EVs isolated from participants with an active infection compared to Controls because the changes in placental pathology and transcriptome were the greatest in AI cases compared to Controls.

To assess the effect of AI EVs on EVT function, we used primary EVTs isolated from first-trimester placenta and quantified EVT invasion through a collagen gel. EVT invasion, which is vital for anchoring the placenta to the uterus and the remodeling of maternal uterine arteries providing blood to the villous trophoblasts, was significantly reduced by exposure to AI EVs compared to Control EVs (Figure 3A). Inadequate invasion and failure to completely remodel maternal arteries increases the risk of preeclampsia, intrauterine growth restriction, and fetal loss (*50, 51*).

**Figure 3.**
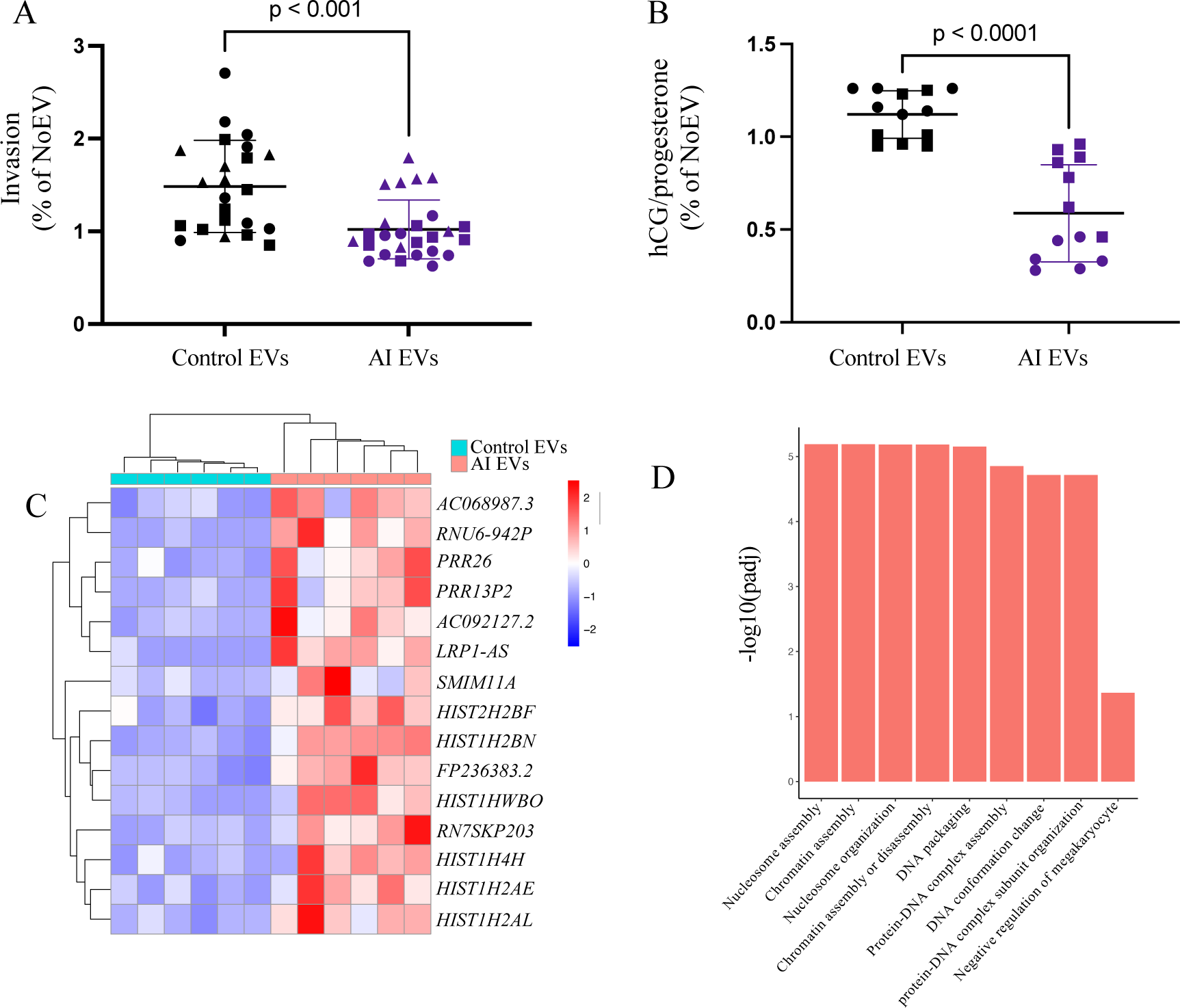
Trophoblast function was disrupted by exposure to EVs isolated from patients with an active infection (AI) compared to Controls. **(A)** Extravillous trophoblasts (EVTs) were isolated from three placentas (identified by symbol shape) and exposed to Control or AI EVs (n=9-10/group, 3 experiments). Invasion was calculated and normalized to invasion of EVTs derived from the same placenta not exposed to EVs (noEVs). All data are presented as mean ± SD. **(B)** Human chorionic gonadotropin (hCG) and progesterone were measured in the media of forskolin-treated (syncytialized) BeWo cells. The ratio of hCG to progesterone was normalized to hormone production by cells not exposed to EVs (noEVs). The results of two experiments (identified by symbol shape) are reported in B. (n=6-7/group, 2 experiments). All analyses were performed by one-way ANOVA or Kruskal-Wallis test, followed by post-hoc tests. **(C)** Following EV exposure, BeWo cell transcriptome was measured, and the top differentially expressed genes comparing the cellular response to AI EVs to Control EVs are listed in the heat map (red represents increased and blue represents decreased expression). **(D)** The biological processes altered by AI EVs compared to Control EVs were determined by Gene Ontology enrichment analysis.

To study syncytiotrophoblasts, we used the BeWo choriocarinoma cell line that is commonly used to study this trophoblast lineage. After the addition of forskolin, BeWo cells syncytialize forming cells that mimic the syncytiotrophoblast, including the production of placental hormones (hCG and progesterone). Syncytiotrophoblast hormone production is essential for the maintenance of pregnancy. The ratio of hCG to progesterone in the media of syncytialized BeWo cells was significantly reduced following exposure to AI EVs compared to Control EVs (Figure 3B). A reduction in the ratio of hCG to progesterone indicates that specific pathways related to steroid hormone production were disrupted. These findings demonstrate that EVs from the circulation of a pregnant individual with an active infection disrupt major trophoblast functions including invasion and hormone production, which may have profound effects on pregnancy maintenance.

To identify novel pathways that may contribute to trophoblast dysfunction, we analyzed the BeWo transcriptome following EV exposure. AI EVs significantly altered gene expression in BeWo cells compared to Control EVs. Multiple genes were dysregulated including genes that encode for long non-coding RNA genes and histone proteins (Figure 3C). This suggests that DNA packaging and transcription is disrupted in trophoblasts exposed to AI EVs compared to Control EVs. Consistent with this, Biological Processes, determined by GO analysis, and top canonical pathways, identified by IPA, were related to cellular transcription and DNA repair (Figure 3D and Supplemental Table 4). The pathways disrupted by AI EVs suggest a generalized effect on trophoblast gene expression that led to disrupted hormone production.

### Increased mtDNA content in LEVs following COVID-19 in early pregnancy

Analysis of the placental transcriptome following COVID-19 identified differentially expressed genes indicative of mitochondrial dysfunction (Table 2). To determine if circulating EVs were enriched in mitochondrial cargo, we measured mitochondrial DNA (mtDNA) content (Figure 4A &C) and found that mtDNA was more abundant in large compared to small EVs. In contrast, nuclear DNA was not consistently measurable in all samples. Additionally, the abundance of mtDNA in large, but not small EVs, inversely correlated with the gestational timing of COVID-19 (Figure 4 B&D). The increase in mtDNA released in large EVs following COVID-19 during early pregnancy suggests that the mitochondrial function of cells producing large EVs was persistently disrupted.

**Figure 4.**
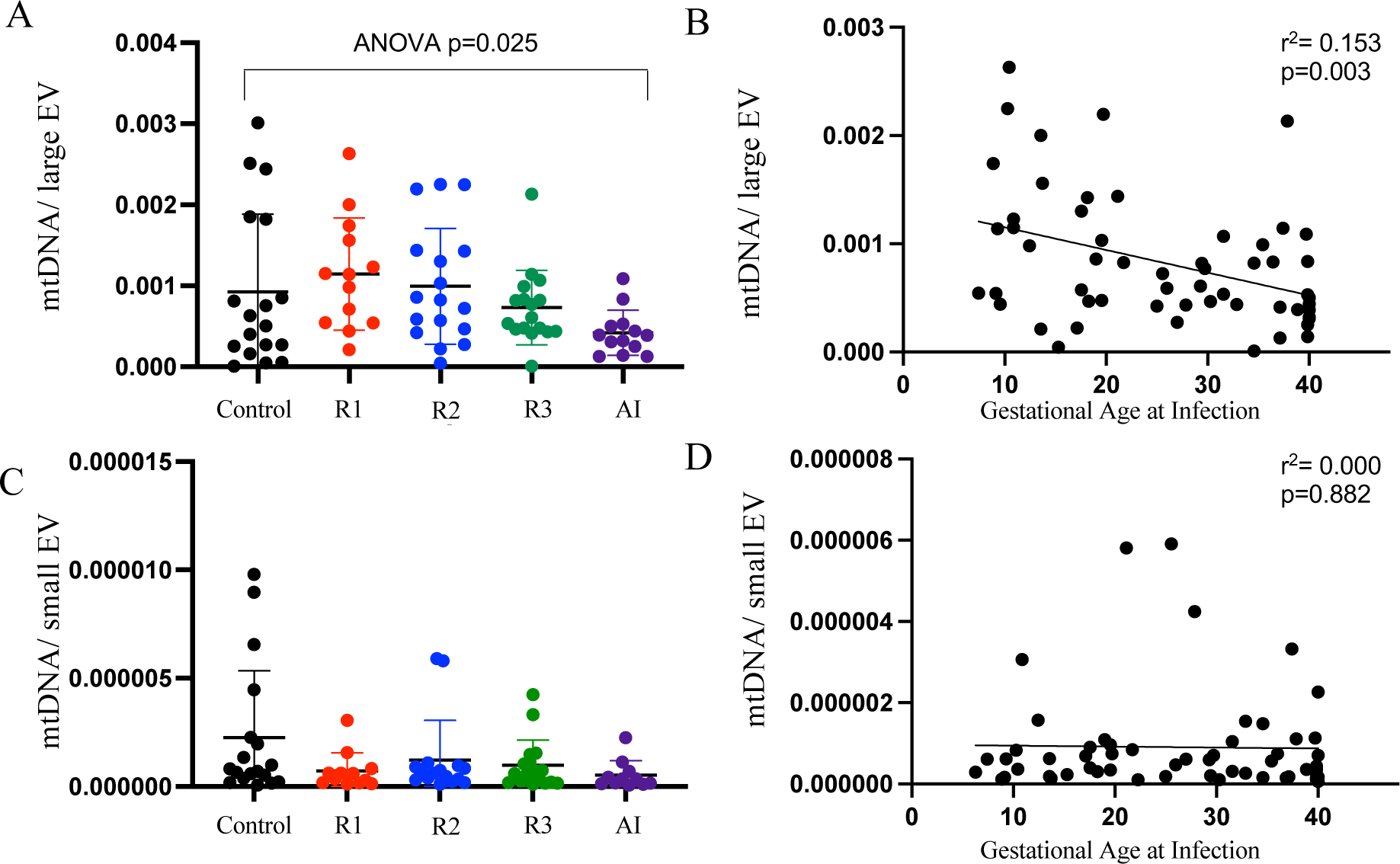
Large EV abundance of mtDNA is inversely correlated with gestational timing of infection. **(A&C)** The amount of mtDNA in each large EV (A) and small EV (C) is reported for each group (n=13-20/group). Data are presented as mean ± SD and tested by ANOVA followed by post-hoc tests. Independent pairwise comparisons were made between Controls and resolved infection in the 1^st^ trimester (R1), second trimester (R2), third trimester (R3), or active infection (AI). **(B&D)** The Pearson correlation between gestational age at infection and mtDNA content is reported for large EV (B) and small EV (D).

**Table 2.**
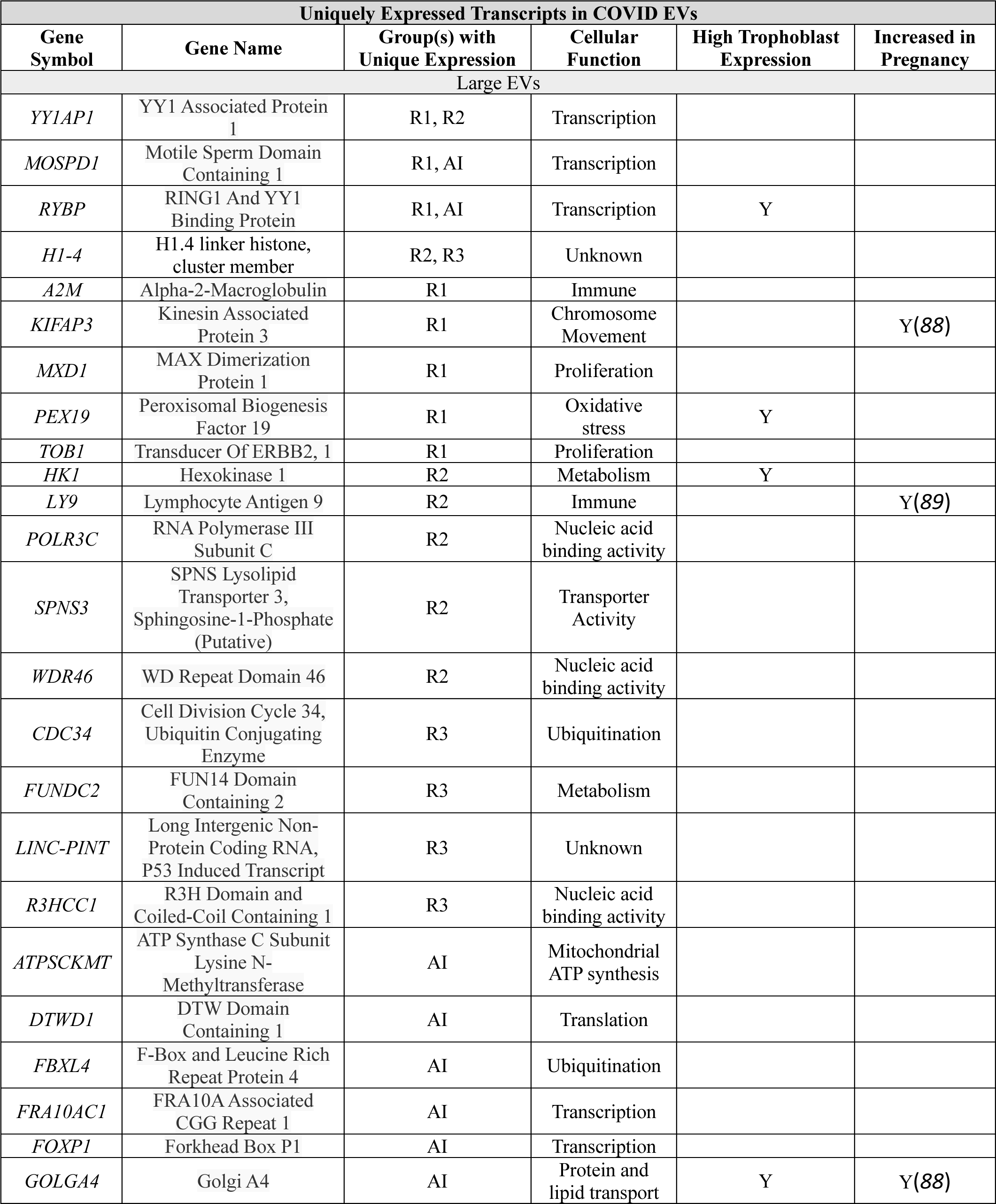
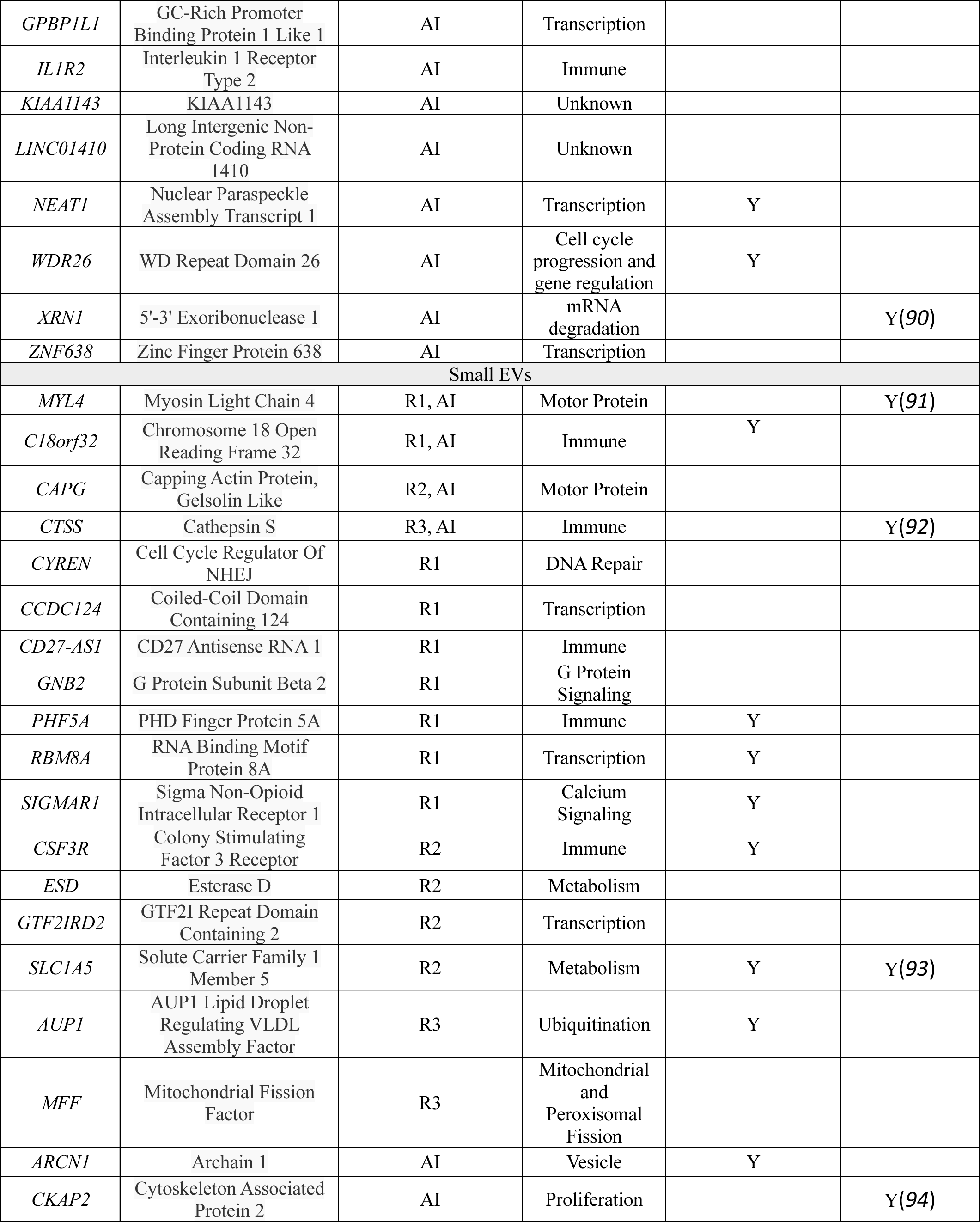
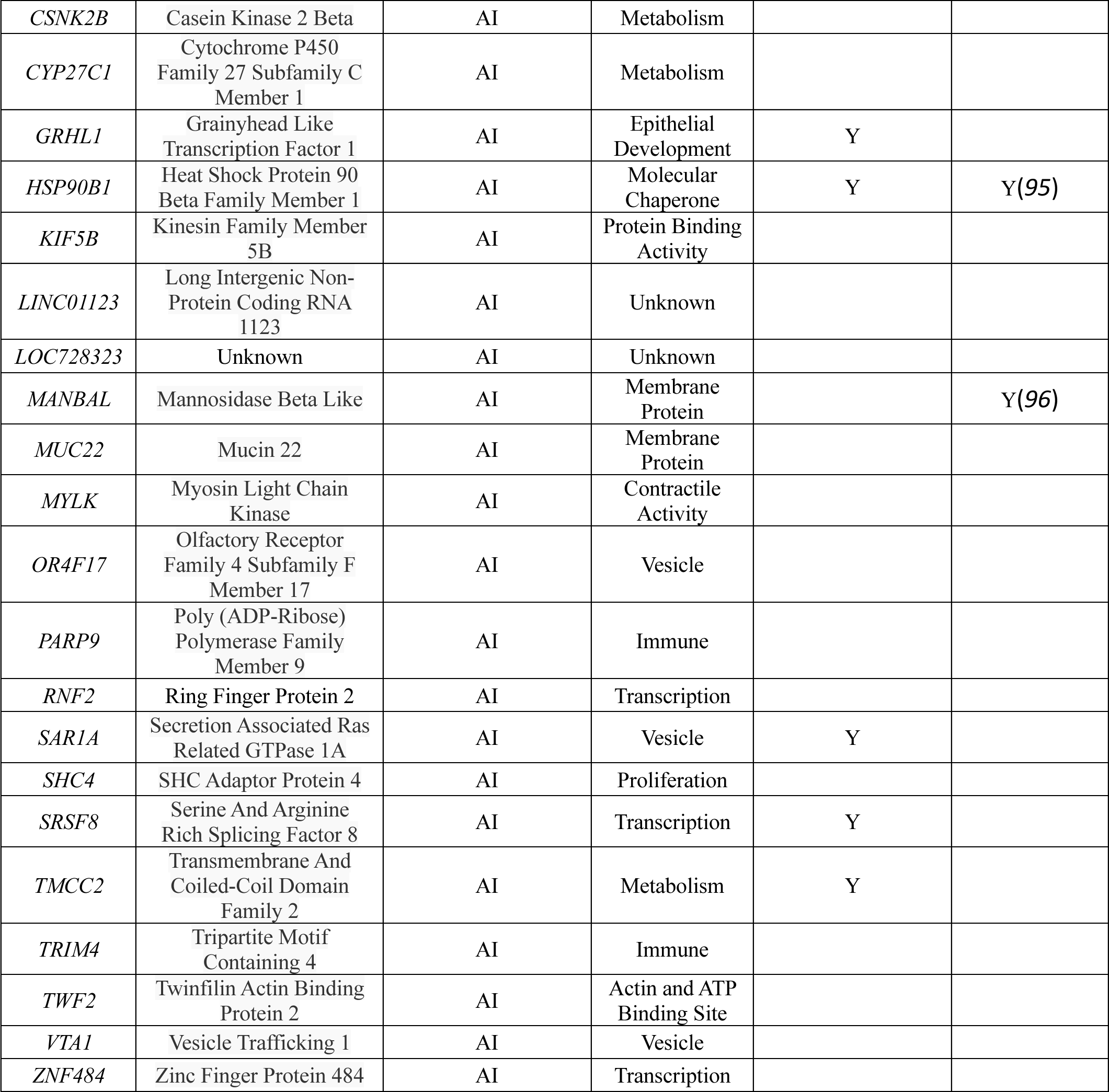
Unique genes identified in EVs isolated from COVID-19 cases Transcripts are carried by large EVs or small EVs isolated from COVID-19 cases that are absent in Controls. The listed transcripts are not detected in EVs isolated from Controls but are present in the identified COVID-19 group(s). The cellular function, expression in trophoblasts, and pregnancy associated expression of each transcript is listed as well (reference listed).

### COVID-19 during pregnancy alters EV RNA cargo

EVs contain small and larger (mRNAs) and long noncoding RNAs, however, small RNAs are the most commonly studied EV cargo (*52*). mRNAs encapsulated within EVs are transferred to recipient cells and translated into proteins, altering the behavior of the recipient cells (*53–56*). Therefore, we profiled the mRNA content of circulating EVs to determine if there were differences dependent on the gestational timing of COVID-19. We sequenced an average of 2,946 gene associated transcripts in large EVs and 1,947 in small EVs.

The most abundant mRNA transcripts in EVs were common to all groups. However, we also identified transcripts that were either uniquely expressed in COVID-19 groups (i.e. absent in Controls), or uniquely expressed in Controls, and absent in one or more COVID-19 groups. Multiple transcripts were uniquely expressed in large EVs from the COVID-19 groups including *YY1AP1*, *MOSPDI, RYBP*, and *H1-4* (Table 2). The proteins encoded by these mRNAs are related to transcription, except for *HI-4* which has an unknown cellular function. In small EVs, *MYL4*, *C18orf32*, *CAPG*, and *CTSS* transcripts were uniquely present in EVs isolated from COVID-19 groups (Table 2). These transcripts encode a motor (*MYL4* (myosin light chain 4)) and immune (*C18orf32* (Putative NF-Kappa-B-Activating Protein 200), *CAPG* (macrophage capping protein), and *CTSS* (cathepsin S)) proteins. This suggests that COVID-19 alters immune-related small EV cargo. Many other transcripts were uniquely detected in EVs isolated from individual COVID-19 groups (Table 2). The unique transcriptome suggests that COVID-19 increased expression of these genes making their transcripts more available for EV packaging or increased specific transcript loading into EVs.

There were several interesting transcripts that were only present in large EVs from Controls, including APBA3, MTSS, FCF1, PSG2, LOC100128233, PHOSPHO2, and THOC3 (Figure 5A). These transcripts encode proteins involved in various cellular functions, including signal transduction, transcription, and proliferation. In contrast, only a few transcripts were unique to Controls in small EVs. These included PYCR2, SCGB1C1, and CD300L4 (Figure 5B). PYCR2 encodes a cellular metabolism protein; CD300L4 encodes an immune-regulating protein. The protein function of SCGB1C1 is unknown. Numerous other transcripts were abundant in Controls but absent in individual COVID-19 groups. If present in the other COVID-19 groups, their expression was decreased compared to Controls (Figure 5).

**Figure 5.**
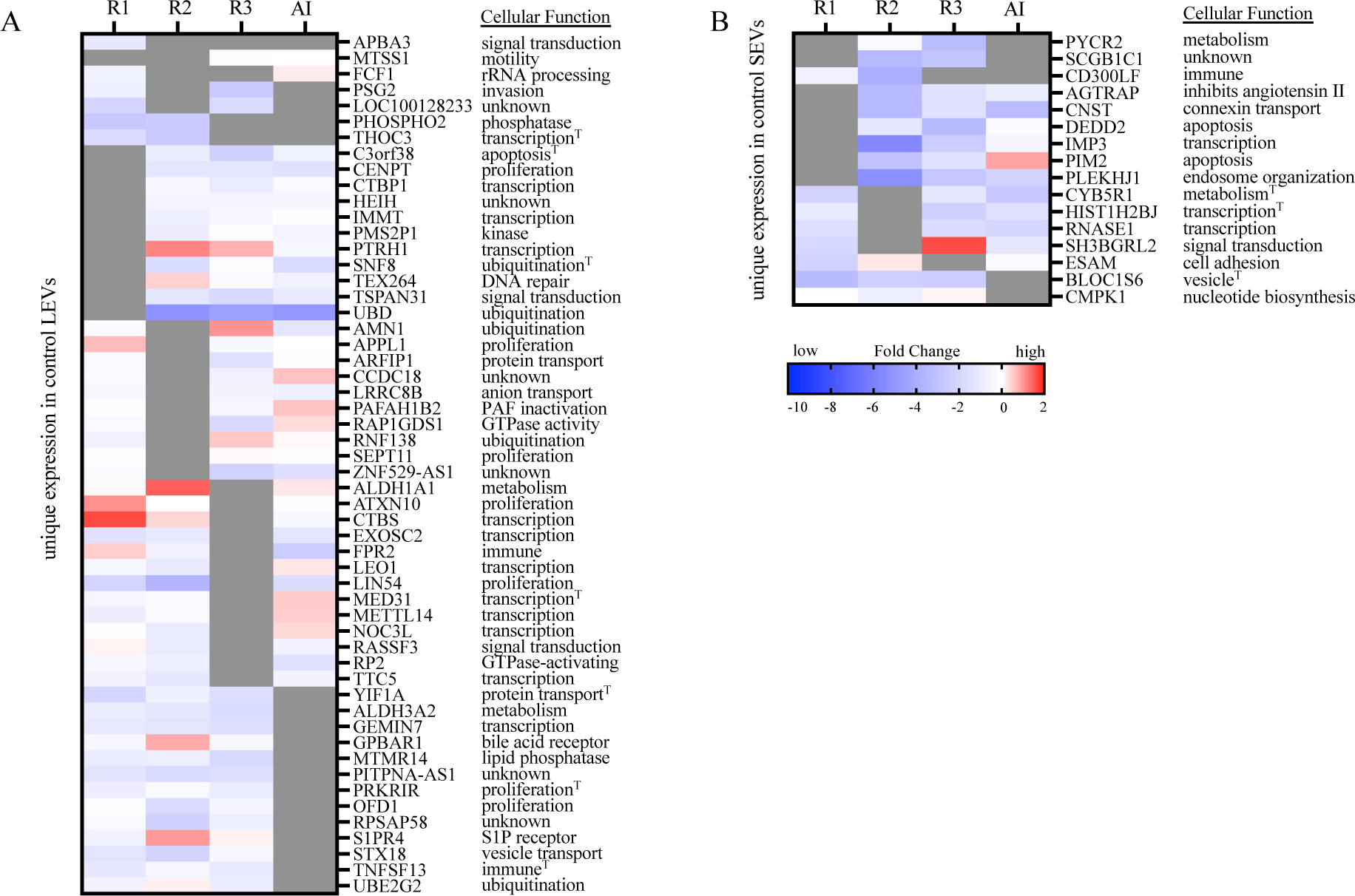
Control EVs carried transcripts that were absent in COVID-19 groups. **(A&B)** mRNA transcripts uniquely detected in EVs isolated from Controls but absent in EVs isolated from COVID-19 groups (gray bars) and differential expression (log 2-fold change) in other COVID-19 groups are listed in the heat map (increased expression is red and decreased expression is blue). The general cellular function of each gene product is listed on the right. Large EV transcripts are reported in (A) and small EV transcripts are reported in (B) (n=9-10/group). Independent pairwise comparisons were made between Controls and resolved infection in the first trimester (R1), second trimester (R2), third trimester (R3), or active infection (AI). Transcripts known to be highly abundant in trophoblasts are marked by (T).

Transcripts carried by EVs reflect the activity of the secreting cell. While the individual transcripts identified in EVs differ, their cellular functions often overlapped (Table 2, Figure 5). For example, large and small EVs isolated from R2 carried transcripts that regulate cell signaling, gene expression, immune regulation, and metabolism. Additional pathways include proliferation, apoptosis, invasion, ubiquitination, platelet function, and vesicle formation.

Many transcripts identified in EVs are abundant in trophoblast cells and have been previously reported to increase with gestation. Interestingly, *THOC3* mRNA, encoded by a highly expressed gene in trophoblast cells, was abundant in Control large EVs but was low or absent in the COVID-19 groups. In addition, large EVs carried a different highly expressed trophoblast transcript, *RYBP*, in COVID-19 groups, but this transcript was absent in Controls. Both proteins encoded by these genes are involved in transcriptional regulation (*57, 58*). Several of the unique mRNAs in EVs isolated from COVID-19 groups have been previously reported to be associated with adverse pregnancy outcomes, including gestational hypertension, preeclampsia, preterm birth, and intrauterine growth restriction (Supplemental Table 5). The number of these pregnancy complication-associated mRNAs was highest in AI EVs, but they were also abundant in EVs isolated from participants with resolved infections. Interestingly, the abundance of eleven transcripts that have been implicated in preeclampsia differed between AI EVs and Control EVs. Of importance, participants with AI also had a higher incidence of preeclampsia. Moreover, two transcripts associated with preterm birth were uniquely carried in R2, but not Control EVs; R2 participants had a higher incidence of preterm birth. This suggests that EV cargo may reflect etiological pathways leading to these pregnancy complications.

### Circulating EVs carry transcriptional regulators of differentially expressed genes in the placenta

Because EVs had a direct functional effect on trophoblasts *in vitro*, we investigated whether they carried transcriptional regulators of genes whose expression was altered in placenta of COVID-19 pregnant participants (Supplemental Table 3). Multiple mRNAs encoding transcriptional regulators were differentially abundant in COVID-19 groups compared to Controls. Several were contained in both small and large EVs (*JUN, FOS, LEPR, LGALS1, CD36*) or only in small EVs *(PRKN*). Moreover, Jun proto-oncogene B (*JUNB*) mRNA levels were increased in large EVs from R1 compared to Controls (FC=2.25, p=0.01). This suggests that *JUNB* carried in EVs could elicit the observed changes in transcription of its downstream targets in the placenta. Similarly, mRNAs encoding 4 of the transcriptional regulators in R3 compared to Control placenta were found in large and small EVs (*IL1B, HIF1A, IGF2, PCBP1*); 1 was only in large EVs (*CXCR4*), and 3 were only in small EVs (*ESR1, AKT1, TP53, SPZ1, IRS2*). Interestingly, interleukin 1 beta (*IL1B*) mRNA was decreased in large EVs from R3 compared to Controls (FC = -1.85, p=0.070), whereas estrogen receptor 1 (ESR1) mRNA levels were lower in R3 small EVs compared to Controls (FC=-6.99, p=0.13). Importantly, expression of genes controlled by these transcriptional regulators was altered in R3 placenta compared to Controls. Similarly, many transcriptional regulators of genes with differential expression in AI compared to Control placenta were found in both large and small EVs (*GRN* and *IL1B*), in large EVs (*FAS*) or in small EVs (*JUN, STAT3, IFG1, IFNG, AGT*). These observations exemplify potential EV-driven signaling leading to altered gene expression in the placenta that occurs in COVID-19.

## DISCUSSION

The long-term effects of COVID-19 during pregnancy have not yet been elucidated. Previous studies and our findings reported here, show that the placenta is damaged, and the likelihood of adverse pregnancy outcomes was increased in patients with a pregnancy complicated by COVID-19. We have begun to elucidate the mechanisms underlying the observed placental abnormalities associated with COVID-19. For the first time, we demonstrate that circulating EVs from COVID-19 affected pregnancies 1) have a detrimental effect on trophoblast function, including hormone production and invasion *in vitro*; 2) are altered after SARS-CoV-2 infection; and 3) carry cargo that has been previously associated with adverse pregnancy outcomes.

Our findings, in *in vitro* experiments, that trophoblast dysfunction following exposure to EVs isolated from study participants with an active SARS-CoV-2 infection provides evidence that circulating EVs contribute to the resulting placental pathology. We focused our *in vitro* trophoblast experiments on the response to EVs isolated from participants with an active infection because the magnitude of alteration in the placental transcriptome was greater compared to those placentas from resolved infections. EVs from patients with an active infection (AI) disrupted fundamental trophoblast functions that are crucial to maintain a healthy pregnancy. Others have shown that trophoblast dysfunction, including failure to invade and produce hormones, contributes to the development of preeclampsia, preterm birth, and intrauterine growth restriction (*50, 51*). Therefore, the AI EV-induced reduction in EVT invasion and syncytiotrophoblast hormone production may have contributed to the development of these pregnancy complications following COVID-19.

Gestational age at the time of infection was a major determinant of COVID-19-induced changes in the profile of circulating EVs. If participants were infected during the first or second trimester of pregnancy, numbers of trophoblast and fetal endothelial cell large EV were increased, and circulating large EVs carried more mtDNA. This suggests that COVID-19 during early pregnancy disrupts mitochondrial function in the placenta. Mitochondrial dysfunction has been reported in many organs following SARS-CoV-2 infection and is thought to contribute to cell injury, cell death, and inflammation (*59–61*). Appelman et al. recently reported persistent mitochondrial dysfunction in skeletal muscle long after the resolution of SARS-CoV-2 infection (*62*). Further, elevated cell-free circulating mtDNA is commonly observed in COVID-19 and correlates with severity and length of infection, reflecting significant mitochondria stress (*63–66*). In support of a direct association of infection and mtDNA release in EVs, Faizan et al. recently demonstrated that SARS-CoV-2 infection causes mitochondrial dysfunction and release of EVs containing mtDNA in airway epithelial cells (*60*). Thus, our results suggest that abnormal mitochondria may also play a role in the pathogenesis of placental dysfunction in COVID-19.

Circulating EV cargo reflects the activity of the cells of origin. The transcripts carried by EVs encode genes related to inflammation, vasculopathies, bioenergetics, and cell death, processes and pathways that were present in the transcriptome and histopathology of the placenta regardless of the timing of infection. EVs carry transcripts that are highly expressed by trophoblasts, and have known functions in cellular metabolism, immune regulation, and transcription. We also found that many of the transcripts in EVs from pregnancies complicated by COVID-19 are encoded by genes that have been implicated in adverse pregnancy outcomes including gestational hypertension, preeclampsia, preterm birth, and intrauterine growth restriction. This points to shared pathways of placental dysfunction induced by a systemic SARS-CoV-2 infection.

EV cargo can elicit a functional response when delivered to a recipient cell, as demonstrated by our *in vitro* studies. While it is not known if mtDNA in EVs per se was responsible for altering trophoblast function in our experiments, multiple studies have demonstrated that mitochondria cargo can alter the recipient cell’s mitochondrial function (*67–69*). mRNA transcripts are also biologically active in recipient cells and we identified transcripts in EVs that encode for multiple transcriptional regulators genes whose expression was altered in placenta following COVID-19. Importantly, expression of several of these genes has been previously reported to be altered in pregnancy complications. For example, JUN signaling was disrupted in R1 placenta compared to Control placenta and JUNB mRNA was increased in Control compared to R1 large EVs. JUN proteins are important for placentation, and a loss of JUN signaling is implicated in preeclampsia (*70, 71*). Thus, low levels of JUNB in COVID-19 EVs may indicate placental dysfunction which in turn could contribute to the later development of preeclampsia, which is observed at higher rates in pregnancies complicated by COVID-19 (*72*). In R3 compared to Controls, hormone receptor signaling was identified as a top canonical pathway and differentially expressed genes were regulated by ESR1. *ESR1* mRNA was abundant in small EVs isolated from Controls but not R3. ESR1 signaling is vital for placental function and pregnancy maintenance because estrogen signaling is obligate for angiogenesis and vasculature control (*73*). In fact, genetic variations in *ESR1* are associated with recurrent pregnancy loss and preeclampsia, and both adverse pregnancy outcomes are increased in maternal SARS-CoV-2 infection during pregnancy (*74, 75*). Thus, our findings suggest placental dysfunction may in fact be a result of EV cargo delivery.

Our study is limited by the number of symptomatic patients with an active infection at the time of delivery. Despite only 14% of pregnant participants experiencing COVID-19 related symptoms, their placentas had significant pathology and an altered transcriptome. This was associated with an increased incidence of preeclampsia and medically indicated preterm birth in asymptomatic and symptomatic AI cases. It’s worth noting that EVs obtained from asymptomatic individuals have been found to exert significant impacts on trophoblast function when tested in vitro. This discovery highlights the importance of exploring the potential consequences of EV exposure in asymptomatic patients and may have important implications for understanding the role of EVs in reproductive health.

Our study has provided significant insights into the profile and functional consequences of circulating extracellular vesicles in mothers who were infected with SARS-CoV-2. This study is the first to demonstrate the negative impact of maternal circulating vesicles on trophoblast function in COVID-19. By comparing the placental transcriptome and EV cargo content, we have identified shared pathways that are associated with pregnancy complications caused by maternal COVID-19 and other pregnancy-related disorders that are not well understood.

## MATERIALS AND METHODS

### Patient cohort

The COMET study was conducted at the Hospital of the University of Pennsylvania (HUP) with Institutional Review Board approval (IRB#843277). Study participants received a description of the study and signed an informed consent before enrollment. Participants were enrolled at the time of delivery in the COMET study between April 2020-June 2022. Participants were tested for a SARS-CoV-2 infection by nasopharyngeal polymerase chain reaction (PCR) upon admission to the labor and delivery unit at HUP. Participants who tested positive at the time of delivery were enrolled in the active infection (AI) group. Those participants who tested negative and had no known SARS-CoV-2 infection during their pregnancy were defined as Controls. Participants with a negative test at delivery and a history of SARS-CoV-2 infection during their pregnancy and greater than 14 days before enrollment, were defined as having a resolved infection (R). All COVID-19 cases were unvaccinated against SARS-CoV-2. The gestational age of SARS-CoV-2 infection was calculated, and participants were further divided into the trimester of infection (resolved infection in the first trimester (R1), resolved infection in the second trimester (R2) and resolved infection in the third trimester (R3)).

### Clinical and demographic data collection

Clinical characteristics, such as maternal age, self-identified race, gestational age at infection, and pregnancy outcomes, were extracted from the medical record (Table 1). The severity of COVID-19 disease was categorized based on the National Institute of Health and Society for Maternal-Fetal Medicine definitions: Asymptomatic infection was defined as participants who tested positive but experienced no symptoms. Symptomatic participants included all levels of illness (mild-critical).

### Sample collection

Placentas were collected at the time of delivery. All placentas were examined by the pathology department at the Hospital of the University of Pennsylvania (HUP). Placentas were assessed using a systematic protocol that includes recording the trimmed placental weight, membrane insertion site, gross appearance, dimensions of the placental disc, and umbilical cord insertion, length, and diameter. Full-thickness placental biopsies were collected from an area devoid of obvious pathology located equidistant between the placental cord insertion and the edge of the placenta. Tissue was fixed in 10% formalin for histological assessment. Macroscopic and microscopic lesions were identified and classified according to the Amsterdam Placental Workshop Group 2014 classification (*76–78*). Placental biopsies were also collected and stored in Trizol for RNA isolation.

Blood was collected at delivery in an EDTA tube and spun at 1,000G for 10 minutes at room temperature to isolate plasma, which was aliquoted and stored at -80°C.

### Placenta and BeWo RNA isolation and sequencing

Total RNA was isolated from placental biopsy samples using Qiagen RNEasy Plus Mini Kits (Cat# 74134 Qiagen, Hilden, Germany). Total RNA was isolated from syncytialized BeWo cells with the PicoPure RNA Isolation Kit (Cat# KIT0204 Applied Biosystems, Waltham, MA). Isolated RNA was sent to NovoGene for library preparation and sequencing.

RNA integrity and quantification were assessed using the RNA Nano 6000 Assay Kit of the Bioanalyzer 2100 System (Agilent Technologies, CA, USA). RNA purity was determined using a NanoPhotometer spectrophotometer (IMPLEN, CA, USA). A total of 1μg RNA per sample was used as input material for the RNA sample preparation. Sequencing libraries were generated using NEBNext Ultra RNA Library Prep Kit for Illumina (NEB, USA) following manufacturer recommendations, and index codes were added to identify samples. Clustering of the index-coded samples was performed on an Illumina Novaseq 6000 sequencer according to the manufacturer’s instructions. After cluster generation, libraries were sequenced, and pair-end reads were generated. Raw data (raw reads) of FASTQ format were processed through fastp. and clean data was obtained by removing reads containing adapter and poly-N sequences and reads with low quality. Pair-end clean reads were aligned to the GRCh38/hg38 reference genome using Spliced Transcripts Alignment to a Reference (STAR) software. FeatureCounts were used to count the read number mapped to each gene. Then RPKM of each gene was calculated based on the length of the gene and read count mapped to the gene. Differential gene expression between COVID-19 groups and Controls was assessed by DESeq2. Differentially expressed genes were determined based on their adjusted p-value (<0.05) and >1.5-fold change. Functional analysis was conducted using Qiagen’s ingenuity pathway analysis (IPA). The clusterProfiler R package was used to perform a Gene Ontology enrichment analysis of genes that were differentially expressed. Canonical pathways, transcriptional regulators, and GO terms were considered significant if the p-value was less than 0.05.

### EV isolation and characterization

Serial centrifugation was utilized to isolate EVs from plasma. One mL of plasma was spun in an Eppendorf 5424 benchtop centrifuge at 2,000 x g for 10 minutes at 4°C. The supernatant was then spun at 20,000 x g for 30 minutes at 4°C. The pellet was washed in 1mL filtered PBS and spun again at 20,000 x g for 30 minutes at 4°C. The large EV pellet was resuspended in 100uL filtered PBS. The supernatant was spun by the Beckman Ultracentrifuge Optima Max TL using the TLA 120.2 rotor at 100,000 x g (48,000RPM) for 90 minutes at 4°C. The pellet was washed with 1mL filtered PBS and spun again at 100,000 x g (48,000RPM) for 90 minutes at 4°C. The small EV pellet was resuspended in 100uL filtered PBS.

EV isolation was confirmed by transmission electron microscopy, nanoparticle tracking, and protein measurement as recommended by the MISEV guidelines (*79*). Transmission electron microscopy images were generated and resulting images reviewed for the presence of EVs. EVs were analyzed by Particle Metrix Zetaview nanoparticle tracking. 11 fields were captured using the following parameters: sensitivity 80, frame rate 30, shutter 100, minimum brightness 1000, minimum area 10, trace length 15. Representative histograms and TEM images for large and small EVs are included in Supplemental Figure 1. CD9 protein abundance was determined by gel electrophoresis. Total protein was measured with a Qubit Protein Assay Kit and EV suspension was evaporated by vacuum and resuspended in electrophoresis buffer. Three large EV (5μg) and small EV (20μg) samples were loaded into BioRad Mini-Protean TGX Gel 4-20% polyacrylamide gels with Licor Chameleon Duo ladders (928–60000) and run at 20mA for 2 hours. Proteins were transferred to nitrocellulose membrane via 200mA over 3 hours on ice. The membrane was blocked in Licor Intercept Blocking Buffer for 1hour at room temperature then incubated with CD9 antibody (HI9a Biolegend Cat 312112) at 1:5000 overnight at room temperature. The membrane was then incubated with Licor IRDye 800CW streptavidin (926-32230) at 1:5000 for 2 hours at room temperature and the membrane was imaged by Licor Odyssey.

### Flow cytometry on EVs

EVs surface protein expression was determined by flow cytometry following the MISEV guidelines (*79*). EVs were resuspended at 1×10^8^/10μL of filtered PBS. Di-8-ANEPPS (Invitrogen Cat# D3167) was reconstituted in ethanol as per manufacturer instruction and further diluted to 1:1000 in filtered PBS. Antibodies were spun at 20,000 x g for 30 minutes at 4°C immediately before use. 10μL of EV suspension was incubated in ANEPPS (10μL) and antibodies, 1.25μL CD45-Ry586 (Cat# BD568135), 1.25μL CD41a-PE/Cy7 (Cat# BDB561424), 1.25μL PLAP-eFlour660 (Fisher Cat# 50-112-4573), and 1.25μL CD34-PE/CF594 (Cat# BDB562449) and 3μL CD31-AF700 (Biolegend Cat# 50-207-2950), for 30 minutes at room temperature. 470μL of filtered PBS was added before samples were measured by BD Symphony A1 cytometer which has improved sensitivity for small particles. Negative controls included: antibodies alone, EVs without Di-8-ANEPPS, and EVs treated with 1% triton. Data was analyzed using FlowJo software. EVs were identified by Small Particle Side Scatter (SP-SSC) and expression of Di-8-ANEPPS and the relative proportion that expresses cell-specific surface proteins was determined by antibody detection.

### *In vitro* trophoblast EV co-culture

Extravillous trophoblasts (EVTs) were isolated from fresh first trimester placenta based on an EVT outgrowth-based protocol established by Gram et al. (*80–84*). In brief, villous tissue was finely minced and cultured at 37°C and 5% CO_2_ in RPMI 1640 media with 20% FBS. After attachment, EVT outgrowth occurs, and those cells were isolated. Isolated EVTs were confirmed by staining for HLA-G and CK7. EVT invasion was measured using the MilliporeSigma Chemicon QCM Collagen Cell Invasion Assay (Cat# ECM558). EVTs were added to the trans-well invasion plate with EV-depleted media and large and small EVs at 1×10^6^/mL. Cells were incubated at 37°C and 5% CO_2_ for 48 hours. Cells that invaded through the collagen membrane were quantified using a fluorescent plate reader (SpectraMax).

BeWo cells, subclone B30, were cultured in 75-cm^2^ flasks (Fisher Scientific) at 37°C and 5% CO_2_ in media (DMEM/F12, 10% FBS, 1% P/S, 1%L-alanyl-L-glutamine). EV-depleted media was made with EV-depleted FBS (Gibco A2720801) and used for cell culture experiments. Cells were plated at 250,000 cells/well in a 6 well plate and 1.5mL of EV-depleted media was added. Cells adhered for 24 hours before adding 1μg/μL forskolin, a cAMP producer, to promote syncytialization for an additional 24 hours. Large and small EVs were resuspended in EV-depleted media at 1×10^6^/mL and added to BeWo cells for an additional 24 hours. At the time of harvest, cell media was collected, and cells released with 0.25% trypsin. Cells were washed and collected as pellets for total DNA measurement and RNA isolation and sequencing. Cell media was spun at 500 x g to clear cell debris and the supernatant was stored for future hormone measurement. Hormones were measured by Penn Fertility Care using Elecsys HCG+β (Cat# 03271749, Roche Diagnostics) and Elecsys Progesterone III (Cat# 07092539, Roche Diagnostics).

### EV mtDNA measurement

mtDNA was isolated and quantified from large and small EVs by TaqMan-based quantitative polymerase chain reaction (qPCR). We determined that 2×10^7^ large EV and 3×10^9^ small EVs were necessary to reliably and robustly measure mtDNA. As previously described, we quantified mitochondrial-encoded human NADH: ubiquinone oxidoreductase core subunit 1 (ND1) as previously described (*85*). The qPCR reactions were performed in triplicates using a QuantStudio 5 Real-time PCR System (Thermo Fisher) using the following thermocycling conditions: 95 °C for 20 s followed by 40 cycles of 95 °C for 1 s, 63 °C for 20 s, and 60 °C for 20 s. Serial dilutions of pooled human placenta DNA quantified for copies of ND1 (copies/µL) by digital PCR (dPCR) were used as a standard curve. The mtDNA amount per EV was determined by normalizing the resulting abundance by the number of EVs in starting material. We calculated the Pearson correlation coefficient to determine the strength of the relationship between gestational age at infection and abundance of EV mtDNA.

### EV mRNA sequencing

Total RNA was isolated from large and small EVs isolated from 500μL of plasma. Isolated EVs were treated with RNAseA (0.02μg/μL) (Invitrogen Cat# 12091021) for 20 minutes at 37°C to degrade extravesicular RNA. Enzyme activity was stopped by freezing samples at -80°C for 5 minutes and immediate resuspension in Trizol. Nucleic acids were isolated via BCP co-incubation, precipitated by isopropanol, and washed in ethanol. mRNA libraries were prepared from total RNA using the SMART-Seq protocol (*86*). Briefly, RNA was reverse transcribed using Superscript II (Invitrogen, Cat#18064014). The cDNA was amplified with 20 cycles and cleaned up with AMPure XP beads (Beckman Coulter Cat#A63881). cDNA was quantified with Qubit dsDNA HS Assay Kit (Life Technologies, Inc. Cat#Q32851), and 2ng of each sample was used to construct a pool of uniquely indexed samples (Illumina Cat# FC-131-1096). A second amplification was performed with 12 cycles and cleaned up with AMPure XP beads. The final library was sequenced on a NextSeq 1000. Data were mapped against the hg19 genome using RSEM and normalized to transcripts per million (tpm)(*87*).

To determine unique expression, we filtered genes to those that had greater than 5 tpm. Unique genes had no expression (≤5 tpm) in all samples in the reference group and had expression (>5 tpm) in the majority (≥ 50% of the samples) in the comparison group (data in Table 2). Expression of unique genes in Controls, but not COVID-19 groups are shown in Figure 4. These strict criteria identified genes in each group that were uniquely expressed, and those genes were considered for subsequent analysis.

### Statistical analysis

Statistical analysis was performed using GraphPad Prism. Differences in participant demographics and outcomes were tested by a chi-squared test and considered statistically different if p<0.05. Data was tested for normality and either parametric or non-parametric tests were used to determine significance. Data points were identified as outliers and removed if they exceeded two times the standard deviation from the mean. A one-way ANOVA tested for a difference within all groups and subsequent post-hoc t-tests or Kruskal-Wallis determined the significance of each COVID-19 group compared to Controls. Pearson’s correlation was used to determine a correlation between gestational age at infection and mtDNA abundance. A p-value less than 0.05 was considered significant. Differential gene expression was determined to be significant if the adjusted p-value was less than 0.05 and the fold change greater than 1.5.

## Acknowledgments

The authors would like to acknowledge the dedication and effort by the Pregnancy and Perinatal Research Center, especially Meaghan McCabe, MPH, in enrolling patients and collecting samples. The authors would also like to thank Dr. Luca Musante at the University of Pennsylvania’s Extracellular Vesicle Core for guidance in EV-related methods. Finally, the authors would like to thank Dr. Jonni Moore and Richard Schretzenmair at the University of Pennsylvania’s Cytomics and Shared Resource Laboratory for their guidance in conducting flow cytometry on EVs.

## Author contributions

Experimental Design: TNG, SM, RLL, RL, LA, MM, CCC, BAK, JFS, SP, RAS

Data acquisition and analysis: TNG, SM, RLL, AW, LA, CCC, BAK

Writing-original draft: TNG, JFS, SP, RAS

Writing-review & editing: TNG, SM, RLL, RL, NAT, AW, LA, MM, CCC, BAK, JFS, SP, RAS

