## Supplemental figure 1 and tables for "Extracellular vesicles alter trophoblast function in pregnancies complicated by COVID-19"

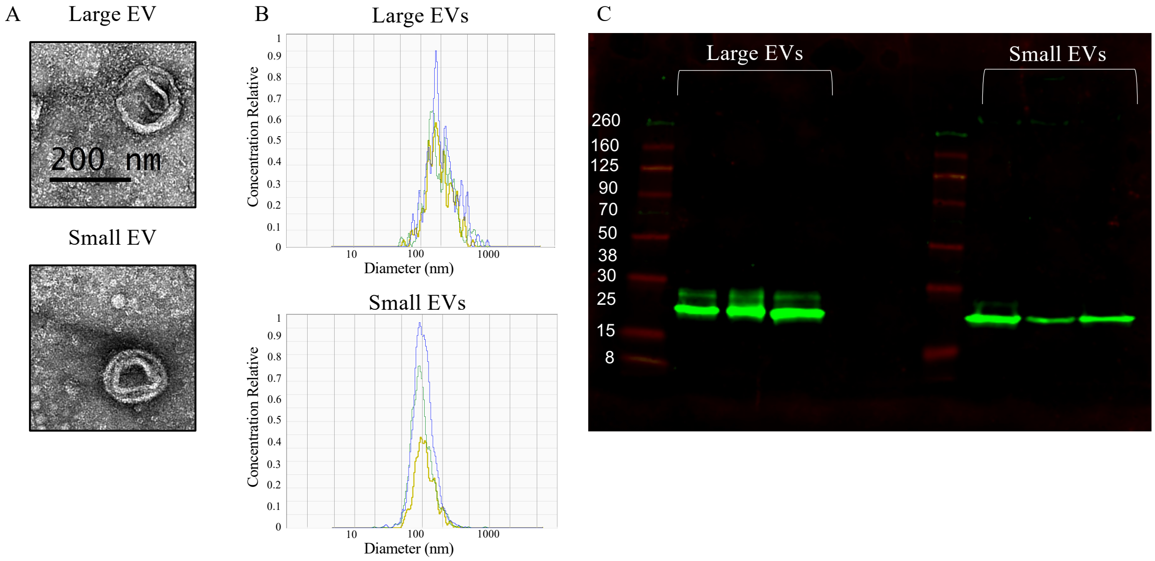


Supplementary Figure 1. Extracellular vesicles isolated from the plasma by serial centrifugation. **(A)** Representative transmission electron microscope images demonstrated EV structure including a lipid membrane. **(B)** Nanotracking analysis was used to determine the size of particles in three Control samples **(C)** CD9 expression evaluated by immune blotting was abundant in isolated particles in three Control samples.

| R1 vs. Control Placenta Transcriptome | | | |
|---|---|---|---|
| **gene symbol** | **gene name** | **FC** | **adj p-value** |
| *GS1-600G8.3* | unknown | 7.48 | 2.09E-04 |
| *AC027288.3* | unknown | 6.85 | 4.35E-04 |
| *RORB* | RAR Related Orphan Receptor B | 6.10 | 3.45E-05 |
| *MMP12* | Matrix Metallopeptidase 12 | 5.20 | 2.22E-04 |
| *OMD* | Osteomodulin | 4.32 | 3.48E-03 |
| *IGFBP6* | Insulin Like Growth Factor Binding Protein 6 | 2.40 | 7.81E-05 |
| *UPK1B* | Uroplakin 1B | 2.59 | 1.16E-05 |
| *FN1* | Fibronectin 1 | 2.38 | 1.41E-06 |
| *SDK1* | Sidekick Cell Adhesion Molecule 1 | 1.62 | 1.15E-04 |
| *MTARC2* | Mitochondrial Amidoxime Reducing Component 2 | -2.15 | 1.67E-05 |
| **R2 vs. Control Placenta Transcriptome** | | | |
| **gene symbol** | **gene name** | **FC** | **adj p-value** |
| *SCN9A* | Sodium Voltage-Gated Channel Alpha Subunit 9 | -1.70 | 0.047 |
| *MAP1LC3C* | Microtubule Associated Protein 1 Light Chain 3 Gamma | -3.37 | 0.008 |
| **R3 vs. Control Placenta Transcriptome** | | | |
| **gene symbol** | **gene name** | **FC** | **adj p-value** |
| *AC079777.1* | Unknown | 6.12 | 4.31E-03 |
| *STAT4* | Signal Transducer and Activator of Transcription 4 | 3.04 | 1.14E-03 |
| *ADARB1* | Adenosine Deaminase RNA Specific B1 | -1.77 | 1.08E-03 |
| *MICAL3* | Microtubule Associated Monooxygenase, Calponin and LIM Domain Containing 3 | -1.97 | 8.89E-06 |
| *SYNPO2* | Synaptopodin 2 | -2.37 | 5.27E-03 |
| *SIGLEC6* | Sialic Acid Binding Ig Like Lectin 6 | -2.46 | 1.37E-04 |
| *PRELP* | Proline And Arginine Rich End Leucine Rich Repeat Protein | -2.52 | 1.64E-03 |
| *ENO2* | Enolase 2 | -3.13 | 4.69E-06 |
| *MEG9* | Maternally Expressed 9 | -3.52 | 4.31E-03 |
| *MAP1LC3C* | Microtubule Associated Protein 1 Light Chain 3 Gamma | -3.80 | 1.37E-04 |
| **AI vs. Control Placenta Transcriptome** | | | |
| **gene symbol** | **gene name** | **FC** | **adj p-value** |
| *PRL* | Prolactin | 6.30 | 6.45E-08 |
| *PENK* | Proenkephalin | 5.10 | 1.84E-05 |
| *TMPRSS3* | Transmembrane Serine Protease 3 | 3.83 | 3.94E-08 |
| *GNLY* | Granulysin | 3.39 | 2.08E-05 |
| *LAMB3* | Laminin Subunit Beta 3 | 2.99 | 2.40E-06 |
| *SLC22A3* | Solute Carrier Family 22 Member 3 | -2.16 | 2.45E-05 |
| *BMP5* | Bone Morphogenetic Protein 5 | -2.56 | 2.42E-05 |
| *CLIC2* | Chloride Intracellular Channel 2 | -2.67 | 2.04E-05 |
| *P2RX1* | Purinergic Receptor P2X 1 | -3.42 | 1.23E-05 |
| *PTGER2* | Prostaglandin E Receptor 2 | -3.45 | 1.33E-05 |

Supplementary Table 1. Placenta RNAseq DEG

Differentially expressed genes (DEG) were identified by comparing COVID-19 groups to controls. The top 10 DEGs are reported with the log 2-fold change (FC) and adjusted p-value.

| R1 v. Control Placenta Transcriptome | | | | |
| --- | --- | --- | --- | --- |
| **canonical pathway** | **FC** | **Z-score** | **p-value** | **# genes** |
| Tumor Microenvironment Pathway | 4.61 | 1 | 0.00 | 4 |
| Osteoarthritis Pathway | 2.83 | nc | 0.00 | 3 |
| Hematopoiesis from Multipotent Stem Cells | 1.94 | nc | 0.01 | 1 |
| IL-17 Signaling | 1.86 | nc | 0.01 | 2 |
| Granulocyte Adhesion and Diapedesis | 1.85 | nc | 0.01 | 2 |
| Hepatic Fibrosis / Hepatic Stellate Cell Activation | 1.83 | nc | 0.01 | 2 |
| Differential Regulation of Cytokine Production in Macrophages and T Helper Cells by IL-17A and IL-17F | 1.76 | nc | 0.02 | 1 |
| Agranulocyte Adhesion and Diapedesis | 1.75 | nc | 0.02 | 2 |
| Pyrimidine Deoxyribonucleotides De Novo Biosynthesis I | 1.66 | nc | 0.02 | 1 |
| Differential Regulation of Cytokine Production in Intestinal Epithelial Cells by IL-17A and IL-17F | 1.66 | nc | 0.02 | 1 |
| **R3 v. Control Placenta Transcriptome** | | | | |
| **canonical pathway** | **FC** | **Z-score** | **p-value** | **# genes** |
| Hepatic Fibrosis / Hepatic Stellate Cell Activation | 2.95 | nc | 0.00 | 4 |
| Tumor Microenvironment Pathway | 2.07 | nc | 0.01 | 3 |
| ILK Signaling | 1.93 | nc | 0.01 | 3 |
| RAR Activation | 1.91 | nc | 0.01 | 3 |
| RHOGDI Signaling | 1.85 | nc | 0.01 | 3 |
| Autophagy | 1.85 | nc | 0.01 | 3 |
| VDR/RXR Activation | 1.84 | nc | 0.01 | 2 |
| Estrogen Receptor Signaling | 1.81 | -1 | 0.02 | 4 |
| TR/RXR Activation | 1.78 | nc | 0.02 | 2 |
| AMPK Signaling | 1.71 | nc | 0.02 | 3 |
| **AI v. Control Placenta Transcriptome** | | | | |
| **canonical pathway** | **FC** | **Z-score** | **p-value** | **# genes** |
| Phagosome Formation | 16.90 | -5.44 | 1.26E-17 | 17 |
| CREB Signaling in Neurons | 11.50 | -3.78 | 3.16E-12 | 26 |
| Hepatic Fibrosis / Hepatic Stellate Cell Activation | 11.00 | nc | 1.00E-11 | 26 |
| GP6 Signaling Pathway | 11.00 | -3.00 | 1.00E-11 | 37 |
| Atherosclerosis Signaling | 10.80 | nc | 1.58E-11 | 41 |
| G-Protein Coupled Receptor Signaling | 9.46 | -3.79 | 3.47E-10 | 19 |
| Breast Cancer Regulation by Stathmin1 | 8.88 | -4.23 | 1.32E-09 | 19 |
| Pulmonary Fibrosis Idiopathic Signaling Pathway | 8.56 | -2.60 | 2.75E-09 | 23 |
| Role of Macrophages, Fibroblasts and Endothelial Cells in Rheumatoid Arthritis | 8.50 | nc | 3.16E-09 | 42 |
| Tumor Microenvironment Pathway | 6.88 | -2.00 | 1.32E-07 | 54 |

Supplementary Table 2. Placenta RNAseq canonical pathways

Ingenuity pathway analysis was used to identify altered pathways in placenta collected from COVID-19 groups compared to control. The top 10 altered pathways are listed for each comparison and the log 2-fold change (FC), z-score, p-value, and number of pathway associated genes are reported. In the event a Z-score is not calculated, nc is reported.

| Transcriptional Regulators of Placenta Transcriptome | | | | |
|---|---|---|---|---|
| **gene symbol** | **gene name** | **activation score** | **p-value** | **# targets** |
| *IGF2* | Insulin Like Growth Factor 2 | 1.96 | 2.41E-05 | 4 |
| *JUN* | Jun proto-oncogene | nc | 4.74E-07 | 7 |
| *UCP3* | Uncoupling Protein 3 | nc | 9.82E-06 | 2 |
| *FOS* | Fos proto-oncogene | nc | 2.01E-05 | 6 |
| *P2RY1* | Purinergic Receptor P2Y1 | nc | 2.06E-05 | 2 |
| *PRLH* | Prolactin Releasing Hormone | nc | 2.06E-05 | 2 |
| *FGF3* | Fibroblast Growth Factor 3 | nc | 3.52E-05 | 2 |
| *LEPR* | Leptin Receptor | nc | 4.13E-05 | 4 |
| *GPX8* | Glutathione Peroxidase 8 (Putative) | nc | 4.40E-05 | 2 |
| *ELN* | Elastin | nc | 6.45E-05 | 2 |
| *LGALS1* | Galectin 1 | nc | 8.82E-05 | 3 |
| *FGF19* | Fibroblast Growth Factor 19 | nc | 9.77E-05 | 3 |
| *PER2* | Period Circadian Regulator 2 | nc | 1.02E-04 | 2 |
| *USP38* | Ubiquitin Specific Peptidase 38 | nc | 1.02E-04 | 2 |
| *FOXC2* | Forkhead Box C2 | nc | 1.04E-04 | 3 |
| *PRKN* | Parkin RBR E3 Ubiquitin Protein Ligase | nc | 1.11E-04 | 3 |
| *RARG* | Retinoic Acid Receptor Gamma | nc | 1.15E-04 | 3 |
| *GAL* | Galanin And GMAP Prepropeptide | nc | 1.33E-04 | 2 |
| *CD36* | CD36 Molecule | nc | 1.60E-04 | 3 |
| *PER1* | Period Circadian Regulator 1 | nc | 2.04E-04 | 2 |
| **Transcriptional Regulators of Placenta Transcriptome** | | | | |
| **gene symbol** | **gene name** | **activation score** | **p-value** | **# targets** |
| *TNF* | Tumor Necrosis Factor | 1.34 | 2.49E-05 | 16 |
| *CYP19A1* | Cytochrome P450 Family 19 Subfamily A Member 1 | 0.882 | 1.52E-04 | 4 |
| *IL1B* | Interleukin 1 Beta | -0.02 | 1.27E-04 | 11 |
| *ESR1* | Estrogen Receptor 1 | -0.27 | 4.36E-05 | 13 |
| *PGR* | Progesterone Receptor | -1.12 | 1.08E-05 | 7 |
| *AKT1* | AKT Serine/Threonine Kinase 1 | -1.42 | 1.03E-05 | 7 |
| *AGT* | Angiotensin | -1.44 | 9,92E-05 | 10 |
| *PPARG* | Peroxisome Proliferator Activated Receptor Gamma | -1.63 | 4.27E-05 | 8 |
| *HIF1A* | Hypoxia Inducible Factor 1 Subunit Alpha | -1.81 | 7.57E-05 | 8 |
| *TP53* | Tumor Protein P53 | -1.9 | 8.31E-05 | 15 |
| *FASLG* | Fas Ligand | nc | 7.57E-06 | 4 |
| *SPZ1* | Spermatogenic Leucine Zipper 1 | nc | 1.94E-05 | 3 |
| *CXCR4* | C-X-C Motif Chemokine Receptor 4 | nc | 2.16E-05 | 4 |
| *PLD1* | Phospholipase D1 | nc | 5.25E-05 | 3 |
| *IGF2* | Insulin Like Growth Factor 2 | nc | 6.81E-05 | 5 |
| *ANGPTL6* | Angiopoietin Like 6 | nc | 9.25E-05 | 2 |
| *MAP2K1* | Mitogen-Activated Protein Kinase 1 | nc | 1.36E-04 | 5 |
| *PCBP1* | Poly(RC) Binding Protein 1 | nc | 1.72E-04 | 2 |
| *IRS2* | Insulin Receptor Substrate 2 | nc | 1.85E-04 | 3 |
| *TRIM29* | Tripartite Motif Containing 29 | nc | 2.21E-04 | 2 |
| **Transcriptional Regulators of Placenta Transcriptome** | | | | |
| **gene symbol** | **gene name** | **activation score** | **p-value** | **# targets** |
| *FAS* | Fas Cell Surface Death Receptor | 1.46 | 5.86E-12 | 33 |
| *GRN* | Granulin Precursor | 1.39 | 3.73E-12 | 19 |

| | | | | |
|---|---|---|---|---|
| *EZH2* | enhancer of zeste 2 polycomb repressive complex 2 subunit | 1.04 | 1.15E-12 | 36 |
| *HRAS* | HRas Proto-Oncogene, GTPase | 0.67 | 3.72E-11 | 45 |
| *AHR* | aryl hydrocarbon receptor | 0.53 | 1.16E-11 | 38 |
| *CEBPB* | CCAAT enhancer binding protein beta | 0.35 | 4.92E-13 | 45 |
| *JUN* | Jun proto-oncogene | 0.04 | 3.54E-12 | 39 |
| *STAT6* | Signal Transducer and Activator of Transcription 6 | -0.07 | 6.20E-14 | 40 |
| *IL13* | Interleukin 13 | -0.91 | 6.95E-19 | 47 |
| *TNF* | Tumor Necrosis Factor | -0.98 | 6.17E-21 | 110 |
| *IL6* | Interleukin 6 | -1.34 | 7.28E-16 | 58 |
| *STAT3* | Signal Transducer and Activator of Transcription 3 | -2.07 | 4.88E-13 | 50 |
| *IL10* | Interleukin 10 | -2.15 | 2.74E-17 | 45 |
| *IL1B* | Interleukin 1 Beta | -2.21 | 5.16E-16 | 71 |
| *IL4* | Interleukin 4 | -2.36 | 6.94E-20 | 81 |
| *IGF1* | Insulin Like Growth Factor 1 | -2.37 | 1.28E-11 | 42 |
| *MAFB* | MAF BZIP Transcription Factor B | -3.02 | 3.53E-11 | 16 |
| *IFNG* | Interferon Gamma | -4.00 | 1.27E-16 | 86 |
| *TGFB1* | Transforming Growth Factor Beta 1 | -4.56 | 2.01E-23 | 115 |
| *AGT* | Angiotensinogen | -4.74 | 1.26E-18 | 67 |

Supplementary Table 3. Placenta RNAseq transcriptional regulators

Ingenuity pathway analysis predicted transcriptional regulators of transcriptional changes in COVID-19 placenta compared to control. The activation score, p-value and number of targets identified in the dataset are listed for the top 20 predicted biological transcriptional regulators.

| BeWo Transcriptome Following Exposure to AI v. Control EVs | | | |
|---|---|---|---|
| **canonical pathway** | **FC** | **p-value** | **# genes** |
| NAD Signaling Pathway | 4.99 | 1.02E-05 | 4 |
| Granzyme A Signaling | 3.86 | 1.38E-04 | 2 |
| DNA Methylation and Transcriptional Repression Signaling | 3.19 | 6.46E-04 | 2 |
| Transcriptional Regulatory Network in Embryonic Stem Cells | 2.95 | 1.12E-03 | 2 |
| NER (Nucleotide Excision Repair, Enhanced Pathway) | 2.39 | 4.07E-03 | 2 |
| Ferroptosis Signaling Pathway | 2.19 | 6.46E-03 | 2 |
| Sirtuin Signaling Pathway | 1.53 | 2.95E-02 | 2 |

Supplementary Table 4. BeWo RNAseq pathway analysis

The canonical pathways disrupted by AI EV exposure compared to Control EV exposure are listed with the log 2-fold change (FC), p-value, and number of pathway associated genes.

| Uniquely Expressed Transcripts Associated with Adverse Pregnancy Outcomes | | | | | |
|---|---|---|---|---|---|
| **Gene Symbol** | **Gene Name** | **EV Population** | **Uniquely Expressed in Group(s)** | **No Expression in Group(s)** | **Adverse Pregnancy Outcome(s)** |
| *CENPT* | Centromere Protein T | LEV | Control | R1 | IUGR(*97*) |
| *IMP3* | IMP U3 Small Nucleolar Ribonucleoprotein 3 | SEV | Control | R1 | PE (*98*) |
| *RAP1GDS1* | Rap1 GTPase-GDP Dissociation Stimulator 1 | LEV | Control | R2 | IUGR (*99*) |
| *LRRC8B* | Leucine Rich Repeat Containing 8 VRAC Subunit B | LEV | Control | R2 | PTB (*100*) |
| *RNF138* | Ring Finger Protein 138 | LEV | Control | R2 | IUGR (*99*) |
| *APPL1* | Adaptor Protein, Phosphotyrosine Interacting with PH Domain And Leucine Zipper 1 | LEV | Control | R2 | IUGR (*101*) |
| *PAFAH1B2* | Platelet Activating Factor Acetylhydrolase 1b Catalytic Subunit 2 | LEV | Control | R2 | PE, gHTN, IUGR (*102, 103*) |
| *CCDC18* | Coiled-Coil Domain Containing 18 | LEV | Control | R2 | PE (*104*) |
| *RNASE1* | Ribonuclease A Family Member 1, Pancreatic | SEV | Control | R2 | PE (*105*) |
| *METTL14* | Methyltransferase 14, N6-Adenosine-Methyltransferase Subunit | LEV | Control | R3 | PE (*106, 107*) |
| *LIN54* | Lin-54 DREAM MuvB Core Complex Component | LEV | Control | R3 | IUGR (*108*) |
| *FPR2* | Formyl Peptide Receptor 2 | LEV | Control | R3 | IUGR (*109*) |
| *ALDH1A1* | Aldehyde Dehydrogenase 1 Family Member A1 | LEV | Control | R3 | IUGR (*110*) |
| *S1PR4* | Sphingosine-1-Phosphate Receptor 4 | LEV | Control | AI | PE (*111*) |
| *THAP12* | THAP Domain Containing 12 | LEV | Control | AI | IUGR (*99*) |
| *TNFSF13* | TNF Superfamily Member 13 | LEV | Control | AI | PE (*112*) |
| *PSG2* | Pregnancy Specific Beta-1-Glycoprotein 2 | LEV | Control | R2, AI | PE, PTB, IUGR (*113-115*) |
| *APBA3* | Amyloid Beta Precursor Protein Binding Family A Member 3 | LEV | Control | R2, R3, AI | PE (*116*) |
| *MXD1* | MAX Dimerization Protein 1 | LEV | R1 | Control | PTB (*117*)<br>PE (*118*) |
| *RYBP* | RING1 And YY1 Binding Protein | LEV | R1 | Control | PE (*57, 119, 120*) |
| *A2M* | Alpha-2-Macroglobulin | LEV | R1 | Control | PE (*121*)<br>PTB (*122*) |
| *HK1* | Hexokinase 1 | LEV | R1 | Control | PE (*123*) |
| *CAPG* | Capping Actin Protein, Gelsolin Like | SEV | R2 | Control | PE (*124*)<br>PTB (*125*) |
| *SLC1A5* | Solute Carrier Family 1 Member 5 | SEV | R2 | Control | PE (*93*) |
| *XRN1* | 5'-3' Exoribonuclease 1 | LEV | AI | Control | PE (*126*) |
| *WDR26* | WD Repeat Domain 26 | LEV | AI | Control | IUGR (*99*) |

| | | | | | |
|---|---|---|---|---|---|
| *GPBP1L1* | GC-Rich Promoter Binding Protein 1 Like 1 | LEV | AI | Control | IUGR (*108*) |
| *FOXP1* | Forkhead Box P1 | LEV | AI | Control | PE (*127*) |
| *NEAT1* | Nuclear Paraspeckle Assembly Transcript 1 | LEV | AI | Control | PE (*128*)<br>IUGR (*129*) |
| *KIF5B* | Kinesin Family Member 5B | SEV | AI | Control | PE (*130*) |
| *CSNk2B* | Casein Kinase 2 Beta | SEV | AI | Control | IUGR (*131*) |
| *MYLK* | Myosin Light Chain Kinase | SEV | AI | Control | PE (*118, 132*) |
| *HSP90B1* | Heat Shock Protein 90 Beta Family Member 1 | SEV | AI | Control | PE (*133*)<br>PTB (*134*) |
| *MUC22* | Mucin 22 | SEV | AI | Control | IUGR (*135*) |
| *GRHL1* | Grainyhead Like Transcription Factor 1 | SEV | AI | Control | IUGR (*136*) |
| *SHC4* | SHC Adaptor Protein 4 | SEV | AI | Control | PE (*137*) |
| *CTSS* | Cathepsin S | SEV | R3, AI | Control | IUGR (*138*) |

Supplementary Table 5. EV Transcripts Associated with Adverse Pregnancy Outcomes

Many transcripts detected in EVs are previously reported to be differentially expressed in adverse pregnancy complications. The gene, EV subpopulation (small [SEV] or large [LEV] EVs), unique expression, and identified literature are listed in the table.
